# Molecular Arms Race: Tannin Biosynthesis and Laccase-Based Detoxification in the Aphid-Gall System

**DOI:** 10.64898/2025.12.15.694272

**Authors:** Qin Lu, Juan Liu, Weiwei Wang, Xin Zhang, Rui He, Gang Cao, Kirst King-Jones, Hang Chen

## Abstract

- Plant-insect coevolution is exemplified by *Schlechtendalia chinensis* inducing gallnuts with record-breaking hydrolyzable tannin (HT) concentrations of 74.49%—32-fold higher than normal leaves. How do plants achieve this extreme defensive chemistry, and how do aphids survive it?
- We integrated transcriptomics, heterologous gene validation in *Arabidopsis*, and enzyme assays to investigate both plant HT biosynthesis and aphid detoxification mechanisms throughout gall development.
- Three key genes—*Phosphoglucomutase* (*PGM*), *UDP-glucosyltransferase BX9* (*BX9*), and *gallate 1-beta-D-glucosyltransferase* (*GDG*) — govern HT biosynthesis, with expression patterns closely tracking tannin accumulation dynamics, and GDG as the rate-limiting enzyme. *Arabidopsis* transformants showed threefold HT increases. Critically, *S. chinensis* employs specialized laccases rather than tannase for detoxification, with laccase activity exceeding tannase by 20,000-fold. The aphid genome encodes three laccase genes, with *Sc-Lac1* expressed in digestive tissues, achieving 39–54% HT degradation.
- These findings reveal how plants weaponize secondary metabolism while herbivores evolve enzymatic countermeasures. The identified genes enable engineering enhanced defenses or pharmaceutical tannin production, while laccase-based detoxification offers new insights into insect adaptation to chemical defenses.

## Introduction

Plant-insect interactions represent one of the most dynamic arenas of coevolutionary innovation, driving reciprocal adaptations that have shaped terrestrial biodiversity over 400 million years (Ehrlich & Raven, 1964; Liu et al., 2023). These interactions encompass a remarkable diversity of strategies, from plants deploying chemical defenses that deter herbivory to insects evolving counter-adaptations that neutralize or even exploit toxic secondary metabolites (Beran & Petschenka, 2022; Berenbaum & Zangerl, 1998). Among the most sophisticated manifestations of this evolutionary arms race are insect-induced plant galls—highly organized structures resulting from parasitic manipulation of host developmental programs to create specialized habitats for the inducing organism (Giron et al., 2016; Lu et al., 2021).

The gall system formed by *Schlechtendalia chinensis* (Hemiptera: Aphididae) on *Rhus chinensis* Mill. (Anacardiaceae) represents an extreme case of metabolic hijacking and serves as a powerful model for investigating plant-insect coevolution. This aphid-plant association produces horned galls containing the highest documented concentrations of hydrolyzable tannins (HT) among plant tissues—reaching up to 74.49% dry weight, more than 30-fold higher than ungalled leaves (Chen et al., 2018; Shao et al., 2013). The commercial significance of these galls, known as Chinese gallnuts, has been recognized for millennia in traditional medicine and industry, yet the molecular basis of this extraordinary HT accumulation remained enigmatic until recent genomic advances. Chromosome-level genome assemblies of both *S. chinensis* (344.59 Mb, 13 chromosomes) and *R. chinensis* (389.40 Mb, 15 chromosomes) have now provided unprecedented insights into this interaction (Ahmad et al., 2024; Ni et al., 2024; Wang et al., 2023). Comparative genomic analyses revealed tandem repeat clusters of key HT biosynthetic genes (*DQD/SDH* and *SCPL* families) specific to HT-rich Anacardiaceae species, alongside transcriptional reprogramming during gall initiation that converts leaf tissue into a highly efficient tannin biosynthesis system (Lu et al., 2025; Ni et al., 2024).

HTs constitute a major class of plant defensive polyphenols restricted primarily to core eudicots, functioning as chemical defenses against herbivores, pathogens, and aluminum toxicity while also providing pharmaceutical benefits to humans (Khanbabaee & van Ree, 2001; Niemetz & Gross, 2005; Okuda et al., 2009). HT biosynthesis initiates with formation of β-glucogallin (1-O-galloyl-β-D-glucose) through conjugation of UDP-glucose and gallic acid via the shikimate pathway, catalyzed by bifunctional dehydroquinate dehydratase/shikimate dehydrogenases (*DQD/SDH*) (Guo et al., 2014; Kasama et al., 2007). Subsequent galloylation steps, mediated by serine carboxypeptidase-like (SCPL) acyltransferases, generate 1,2,3,4,6-penta-O-galloyl-β-D-glucose (PGG), the pivotal intermediate for complex gallotannins and ellagitannins (Grundhöfer et al., 2001; Ono et al., 2012). Recent functional characterization has identified UDP-glucose: gallate glucosyltransferases (UGT84A family) as rate-limiting enzymes whose expression is stringently regulated by *WRKY* and *HD-Zip* transcription factors (Huang et al., 2015; Ishida et al., 2022; Mittasch et al., 2014). In oak species (*Quercus*), differential expression of *UGT84A13* and polymorphisms in its promoter region drive interspecific variation in HT accumulation (Ishida et al., 2022; Mittasch et al., 2014). Moreover, metabolic engineering approaches have successfully reconstituted HT biosynthesis in non-accumulating plants like *Nicotiana benthamiana*, demonstrating the modular and transferable nature of this pathway (Liu et al., 2013). While the general HT biosynthetic framework is established, critical questions remain regarding how gall-inducing aphids induce such extreme (>30-fold) HT elevation: (i) Which host genes are specifically upregulated? (ii) What are their temporal dynamics during gall ontogeny? (iii) How do aphid effectors orchestrate this metabolic transformation? The ability of *S. chinensis* to thrive in this extreme high-tannin environment poses an equally compelling evolutionary puzzle. Tannins impose severe physiological constraints on herbivorous insects through protein precipitation (reducing digestibility), chelation of essential nutrients (iron, zinc), generation of reactive oxygen species, and direct toxicity to gut epithelia (Barbehenn & Peter Constabel, 2011; Hagerman & Butler, 1991; Salminen & Karonen, 2011). Specialized tannin-adapted insects have evolved diverse counter-strategies, including behavioral avoidance, gut pH modulation, production of tannin-binding salivary proteins, and enzymatic detoxification systems (Barbehenn et al., 2006). Among enzymatic adaptations, two principal enzyme classes mediate tannin metabolism: tannases (tannin acyl hydrolases, EC 3.1.1.20) that hydrolyze ester and depside bonds in gallotannins, and laccases (benzenediol: oxygen oxidoreductases, EC 1.10.3.2) that catalyze oxidative transformations of phenolic compounds (Aguilar et al., 2007; Dwivedi et al., 2011). While tannase-based detoxification is well-characterized in certain beetle and moth lineages feeding on tannin-rich oak foliage (Konno et al., 1999; Schuler & Berenbaum, 2013), laccase-mediated degradation of plant secondary metabolites represents an emerging paradigm in insect-plant interactions. Laccases function through free radical generation mechanisms and have been implicated in lignin degradation by wood-boring insects and detoxification of plant phenolics in various herbivore systems (Dittmer & Kanost, 2010; Peng et al., 2018). However, the relative contribution of tannase versus laccase systems in aphids colonizing ultra-high-tannin environments remains unexplored. Genome-wide analysis of *S. chinensis* identified extensive detoxification gene repertoires, including 38 cytochrome P450s, 24 glutathione-S-transferases, and 18 carboxylesterases, suggesting sophisticated xenobiotic metabolism capacity (He et al., 2022). Yet whether HT detoxification relies on canonical hydrolytic mechanisms or alternative oxidative pathways has not been experimentally determined.

This study addresses fundamental gaps in understanding the *S. chinensis*-*R. chinensis* interaction cascade through integrated molecular, biochemical, and functional genomic approaches. Our objectives are threefold: (1) identify and functionally characterize the core regulatory genes governing HT biosynthesis in gall tissue, with emphasis on their expression dynamics throughout gall development; (2) elucidate the enzymatic basis of *S. chinensis* tolerance to extreme HT concentrations, specifically testing the hypothesis of laccase-mediated versus tannase-mediated detoxification; and (3) construct an integrative model of the "aphid effector-induced stimulation → plant biosynthetic reprogramming → aphid enzymatic adaptation" cascade. By combining gene expression profiling, heterologous functional validation, and comparative enzymatic assays, we seek to illuminate both sides of this coevolutionary arms race—revealing how aphids orchestrate host metabolic transformation while simultaneously evolving biochemical mechanisms to tolerate the very toxins they induce.

## Materials and Methods

### Materials

The aphids *S. chinensis* were cultivated on *R. chinensis* in the greenhouse of the Research Institute of Resource Insects of Kunming, Yunnan Province, China (25°02′14″ N, 102°42′33″ E). Horned gall samples were collected at 30 days post-development, with subsequent collections conducted at 60, 70, 80, 90, 100, 120, 145, and 165 days of growth. Upon harvest, aphids within the galls were carefully removed using a soft brush. The gall wall tissues were divided into three groups: the first group was separated into inner and outer layers and air-dried at room temperature; the second group was air-dried for subsequent analysis of HT content; and the third group was immediately frozen in liquid nitrogen for real-time quantitative PCR and gene cloning.

Aphid samples from each developmental stage were also divided into three portions: the first portion was homogenized immediately for assays of tannase and laccase activities; the second portion was homogenized and transferred into dialysis bags (molecular weight cut-off: 10–12 kDa) containing 1× PBS buffer (supplemented with 1% BSA), which were then immersed in fresh buffer on ice for 20 min with gentle stirring to remove small molecular weight compounds, after which the resulting solution was collected as crude enzyme extract for tannase activity measurement; the third portion was frozen in liquid nitrogen for subsequent use in real-time quantitative PCR. Furthermore, aphids collected at 80 days of development were dissected under a stereomicroscope to separate heads from abdomens. The isolated tissues were immediately homogenized in 1× PBS buffer containing 1% BSA on ice for laccase activity measurement. Ensiform galls derived from *Rhus potaninii* were collected in Emei City, Sichuan Province, China (29°36′N, 103°29′E).

The HT content in gall tissues was quantitatively analyzed using the Tannic Acid Content Assay Kit (Colorimetric Method, BioVision, San Francisco, CA, USA). The activities of laccase and tannase in aphids were determined using the Laccase Activity Assay Kit (Micromethod, BioVision, San Francisco, CA, USA) and the Tannase Activity Assay Kit (Micromethod, BioVision, San Francisco, CA, USA), respectively. All assays were performed in triplicate with independent biological replicates to ensure reproducibility.

### Validation of Transcriptomic Data via real-time quantitative PCR (RT-qPCR)

Primers were designed using Primer Premier 6.0 (Table S1) based on transcriptome data (Premier Biosoft International, 2007). Total RNA from plant tissues (gall walls) and aphids was extracted using the TaKaRa MiniBEST Plant RNA Extraction Kit (TaKaRa, Dalian, China) and the TaKaRa MiniBEST Universal RNA Extraction Kit (TaKaRa, Dalian, China), respectively, according to the manufacturer’s instructions. First-strand cDNA was synthesized using the PrimeScript™ RT reagent Kit with gDNA Eraser (Perfect Real Time) (TaKaRa, Dalian, China). Real-time quantitative PCR was performed using SYBR Premix Ex Taq (Perfect Real Time) (TaKaRa, Dalian, China) on a CFX96 Touch Real-Time PCR Detection System (Bio-Rad, Hercules, CA, USA).

For plant gene expression analysis, β*-actin* was selected as the reference gene, and *R. chinensis* leaves were used as the control. For aphid gene expression analysis, β*-actin* was also selected as the reference gene, and overwintering nymphs living on moss were used as the control. The relative expression levels were calculated using the 2^(-ΔΔCt) method. All RT-qPCR reactions were performed in triplicate with three independent biological replicates.

### Gene Cloning

Young *R. chinensis* leaves were collected and homogenized in a liquid nitrogen bath. Total RNA was extracted from 50 mg tissue using an RNA prep pure Tissue kit (TianGen, Dalian, China), and RNA purity and concentration were assessed using a spectrophotometer (NanoDrop 2000, Waltham, MA, USA). A SMARTer Kit (Clontech, USA) was used for 5′- and 3′-rapid amplification of cDNA ends (RACE) using a nested PCR procedure with gene-specific primers and nested gene-specific primers (Table S2, Table S3). The first round of 5′- and 3′-RACE amplification was performed with UPM (Universal Primer A Mix, Clontech, USA) and GSP according to the manufacturer’s protocol. The products from the first round of amplification were used as templates for nested PCR reactions using the nested primers NUP (Nested Universal Primer A, Clontech, USA) and NGSP. All nested PCR reactions were performed in 50 µL reaction mixtures containing 2.5 µL of template, using Ex Taq HS (TaKaRa, China) (Table S3). The nested PCR annealing temperatures for the 3′ and 5′ sequences were 58°C and 68°C, respectively, with all other conditions following the kit manufacturer’s instructions.

Amplified PCR products were analyzed using 1.5% agarose gel electrophoresis and extracted using a SanPrep DNA Gel Extraction Kit (Sangon Biotech, China). The pEASY-T5 zero vectors (TransGen, China) were used to clone target genes, and the constructs were transformed into trans1-T1 cells (TransGen, China). Six plasmids each from 5′- and 3′-RACE were randomly selected and sequenced. The genes’ ORFs and peptide sequences were identified using the NCBI ORF Finder (https://www.ncbi.nlm.nih.gov/orffinder/).

Multiple sequence alignments of PGM, BX9, and GDG amino acid sequences were performed using MEGA (v15.11) with the MUSCLE algorithm. Homologous sequences from related Anacardiaceae and Sapindales species (*Pistacia vera*, *Mangifera indica*, *Citrus × changshan*, *Citrus sinensis*, and *Melia azedarach*) were retrieved from NCBI GenBank. Conservation analysis was conducted using the built-in conservation scoring function in MEGA, which evaluates amino acid identity and similarity across aligned sequences. Residues were classified into conservation categories based on percentage identity: completely conserved (100% identical across all species), highly conserved (strong conservation with minimal variation), moderately conserved (conservation of physicochemical properties), and variable (frequent substitutions). Alignment visualization was performed using default MEGA coloring scheme, where background colors reflect conservation levels (Tamura et al., 2021). The amino acid sequences were submitted to the SWISS-MODEL protein structure prediction server (https://swissmodel.expasy.org/interactive) for structural modeling (Waterhouse et al., 2018). The resulting .pdb files were imported into PyMOL (v3.1) for refinement and structural visualization (Schrödinger, 2015). Secondary structural elements, including α-helices, β-sheets, β-turns, and loops/random coils, were identified and quantified based on the predicted models.

### Gene Overexpression

Two pairs of gRNA primers were designed based on the gene sequences using the CRISPOR web tool (http://crispor.tefor.net/)(Concordet & Haeussler, 2018). The specificity of the gRNAs was validated using an online tool provided by the UCSC Genome Browser (http://genome.ucsc.edu/) (Kent et al., 2002). Following confirmation of primer accuracy, sticky ends (CACC, CAAA) were incorporated, and 6×His and Flag tag sequences were introduced into the downstream primer (Table S4). The primers were synthesized by a commercial biotechnology company.

The target fragments were amplified using Phanta Max Super-Fidelity DNA Polymerase (Vazyme). The pCambia1301 vector was linearized using BglII and BstEII double digestion. Subsequently, the purified digestion products were recombined with cloned PCR fragments of the *PGM*, *BX9*, and *GDG* genes using Vazyme’s ClonExpress-II One Step Cloning Kit. The recombinant products were transformed into *E. coli* competent cells. Positive transformant colonies were detected by PCR, and the PCR products were sequenced for confirmation before plasmid extraction.

The previously constructed single-gene expression vector plasmids were linearized individually using KpnI. Using the expression vectors containing the target gene fused to 3×Flag as templates, primers were designed to amplify the target gene + 3×Flag fragment for the construction of dual-gene expression vectors (*PGM*_*GDG*, *PGM*_*BX9*, *BX9*_*GDG*). Following purification of the restriction products, recombination reactions were performed with the amplified target gene fused to 3×Flag. The resulting products were transformed into *E. coli* competent cells. Positive transformant colonies were identified by PCR, confirmed through sequencing of PCR products, and plasmids were extracted.

Transformed plasmids were introduced into *Arabidopsis thaliana* via Agrobacterium-mediated transformation, with 10 plants infected per gene construct. At day 50 of cultivation, individual plant weight, height, leaf length, and width were measured, followed by immediate rapid freezing in liquid nitrogen for storage. Target gene expression levels were quantified using real-time fluorescent quantitative PCR. The three plants with the highest expression levels in each treatment group were subjected to Western blot analysis to detect target protein expression. UDP-glucose and HT content were determined by liquid chromatography.

### Statistical Analysis

Data were presented as mean ± standard error (SE) from three independent biological replicates, each with three technical replicates. Differences between treatments were analyzed using one-way ANOVA followed by Fisher’s LSD post-hoc test (α = 0.05) in SAS 9.1 (Inc, 2006). For gene expression analysis, fold changes >2 with P < 0.05 were considered significant.

## Result

The gall induced by the fundatrix of the aphid *S. chinensis* on its host plant *R. chinensis* contains a substantial amount of water-soluble tannic acid, with glucose serving as the primary biosynthetic precursor. Glucose is subject to a series of enzymatic transformations that ultimately result in its conversion into tannin, a structurally complex substance. Preliminary transcriptomic analyses have identified three key regulatory genes involved in tannic acid biosynthesis: *Phosphoglucomutase* (*PGM*), *UDP-glucosyltransferase BX9* (*BX9*), and *gallate 1-beta-D-glucosyltransferases* (*GDG*), *PGM* regulates upstream steps in the biosynthetic pathway, whereas *BX9* and *GDG* function in downstream enzymatic processes (Fig. 1A). During the period of gall development, there is a significant fluctuation in HT content, which remains at a comparatively higher level than that observed in leaves. At the initial stage of gall formation, HT levels are recorded at 21.13%. As aphid-induced stimulation progresses, the concentration in the gall wall increases steadily, reaching a maximum of 74.49% on day 80 post-gall induction. Subsequent to this, a gradual decline is observed, with levels decreasing to 60.79% at gall dehiscence. In contrast, the HT content of *R. chinensis* leaves exhibits a consistent low level, ranging between 2.24% and 3.75%. This finding suggests that gall HT concentrations can be approximately 32-fold higher than those found in leaves (Fig. 1B). Real-time quantitative PCR analysis reveals a close correlation between the expression patterns of the key regulatory genes and HT accumulation. It is noteworthy that *GDG*, the terminal gene in the pathway, exhibits the highest expression level (Fig. 1C).

**Figure 1.**
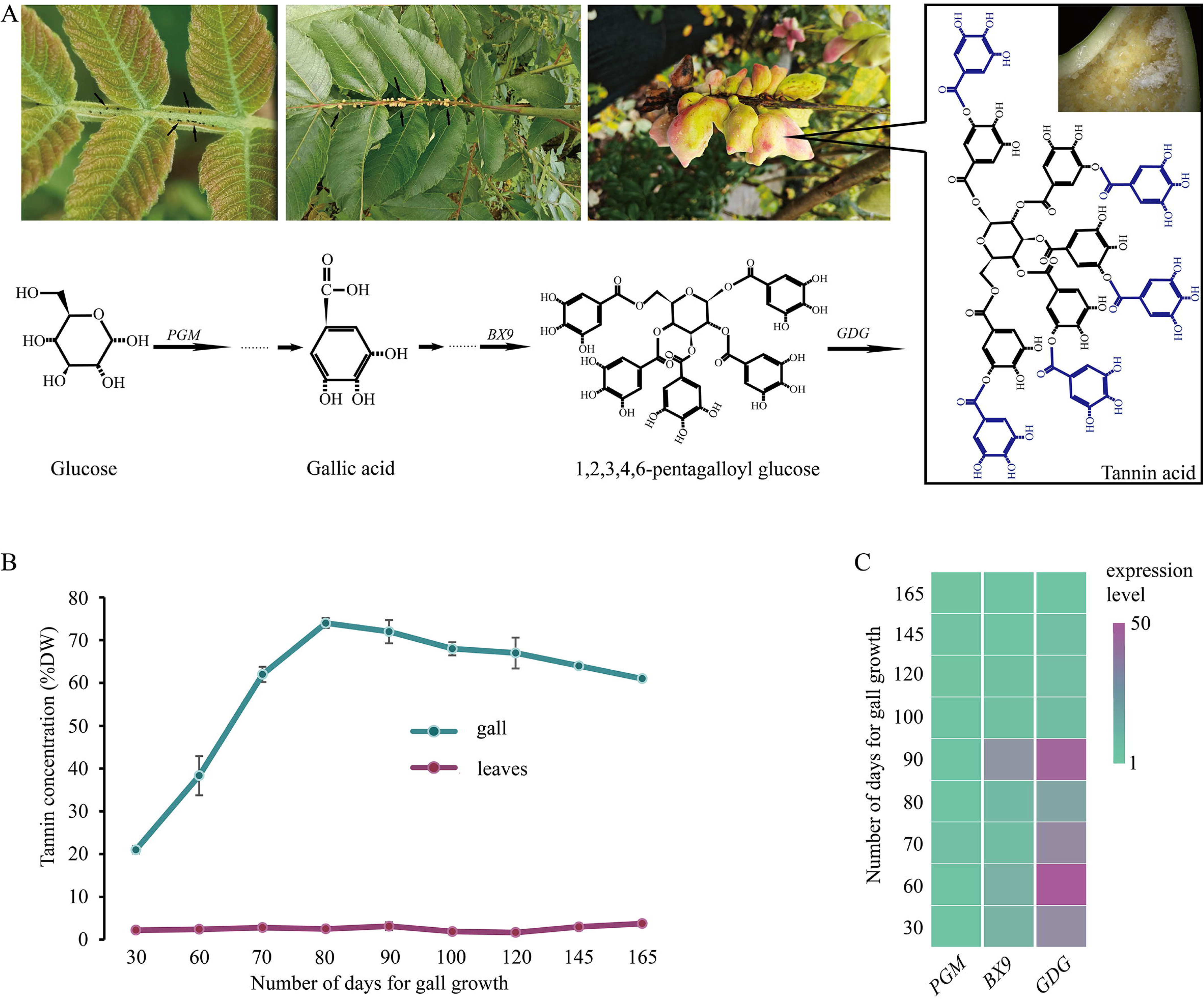
HT biosynthesis dynamics and correlation with gene expression during gall development. **(A)** Schematic representation of the HT biosynthesis pathway showing the regulatory roles of *PGM*, *BX9*, and *GDG*. PGM regulates upstream steps in the biosynthetic pathway, converting glucose-6-phosphate to glucose-1-phosphate. *BX9* and *GDG* function in downstream enzymatic processes, with *BX9* catalyzing UDP-glucose formation and *GDG* catalyzing the final steps in gallotannin biosynthesis. **(B)** Temporal dynamics of HT accumulation in gall walls and leaves throughout gall development. During gall development, HT content exhibits significant fluctuation and remains at comparatively higher levels than observed in leaves. **(C)** Real-time quantitative PCR analysis of key regulatory gene expression patterns during gall development. Expression levels show close correlation with HT accumulation dynamics.

After gall formation, aphids establish feeding sites within the enclosed structure. Stylet penetration into the inner gall wall leaves characteristic stylet sheaths, creating slightly darkened puncture zones (Fig. 2A-D). Notably, despite concentrated feeding activity on the inner wall, HT content had no significant differences between inner and outer walls throughout the growth period (Fig. 2E). Aphids inevitably suck up tannin (Rodríguez et al., 2022); nevertheless, tannase activity within the insects remains at a remarkably low level, which only approximately 0.08 nmol/min/g during the late developmental stages. On the other hand, laccase activity remains consistently elevated across aphid generations within the gall, reaching a peak of 1649.86 nmol/min/g during the early developmental stage (30 days post-induction) and remaining above 1294.57 nmol/min/g until day 90 of gall development. In the later stages of development, laccase activity declines progressively, reaching a nadir of 300.01 nmol/min/g at the last stage. Sexual aphids inhabiting moss where HT is not consumed have been observed to exhibit lower laccase activity, measured at 127.47 nmol/min/g (Fig. 2F). A marked difference in laccase activity is also observed between the head and abdomen of the fundatrix, the activity in the head is 83.04 nmol/min/g, whereas in the abdomen it reaches 886.28 nmol/min/g, indicating that laccase is predominantly localized in the abdominal segments (Fig. 2G).

**Figure 2.**
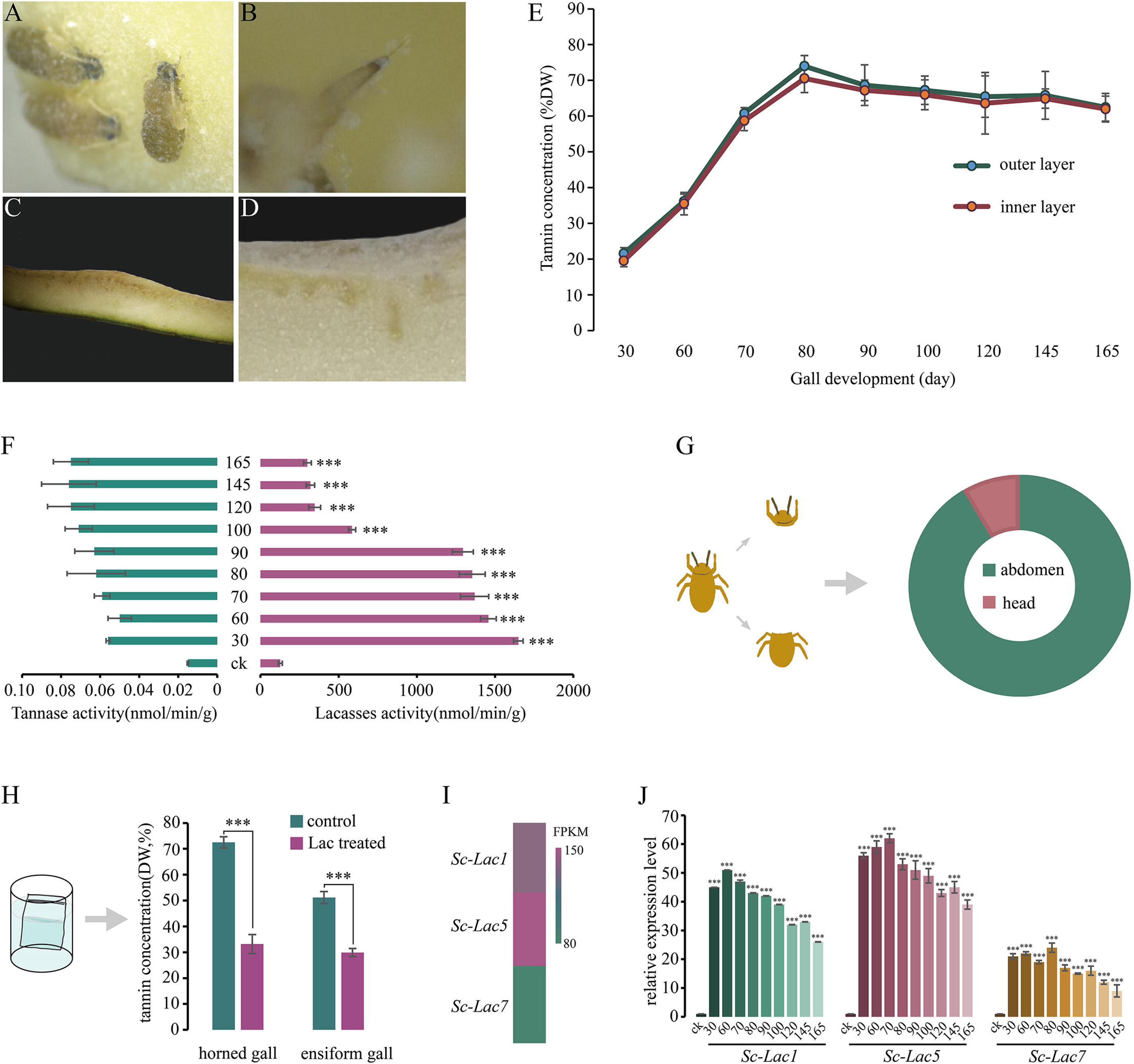
Laccase activity, gene expression, and gall tannin dynamics in aphid-induced galls. **(A-D)** Microscopic examination of stylet penetration sites on the inner gall wall. **(E)** Comparison of HT content between inner and outer gall walls throughout gall development. Despite concentrated aphid feeding activity on the inner wall, no significant differences in HT content were observed between inner and outer walls during the growth period. **(F)** Dynamic changes in tannase and laccase activities across different aphid developmental stages. **(G)** Tissue-specific distribution of laccase activity in the fundatrix. **(H)** Effect of aphid homogenate treatment on HT levels in gall-derived solutions. **(I)** Relative expression levels of three laccase-encoding genes (*Sc-Lac1, Sc-Lac5,* and *Sc-Lac7*) identified in the aphid genome. *Sc-Lac1* exhibits the highest expression level, followed by *Sc-Lac5,* while *Sc-Lac7* shows the lowest expression. **(J)** Temporal expression patterns of laccase genes during gall development. * = P < 0.05; ** = P < 0.01; *** = P < 0.001.

Two types of gall-derived solution were incubated with aphid homogenate. Following treatment, a significant reduction in HT levels was observed in both solutions. In the horned gall solution group, content decreased from 72.52% to 33.19%, and in the ensiform gall group, it declined from 51.19% to 29.91% (Fig. 2H). The aphid genome contains three laccase-encoding genes: *laccase 1* (*Sc-Lac1*), *laccase 5* (*Sc-Lac5*), and *laccase 7* (*Sc-Lac7*). Among these, *Sc-Lac1* exhibits the highest expression level, followed by *Sc-Lac5*, while *Sc-Lac7* shows the lowest (Fig. 2I). Real-time quantitative PCR analysis further confirmed that the expression dynamics of these genes align with transcriptomic data. The highest level of *Sc-Lac1* is observed on day 60 of gall development, the *Sc-Lac5* peak on day 70, and the *Sc-Lac7* peak on day 80. The expression of all three genes gradually declines as gall development proceeds and remains significantly higher than in the control group (Fig. 2J).

The full-length genes of *PGM*, *BX9*, and *GDG* are 1,631 bp, 1,786 bp, and 1,712 bp, respectively. Sequence analysis revealed that each gene encodes multiple polypeptide chains with distinct functional domains. *PGM* encodes a 38-amino acid peptide (98–184 bp) and a larger 312-amino acid polypeptide (438–1376 bp). Similarly, *BX9* produces a 59-amino acid N-terminal peptide (8–187 bp) and a 465-amino acid catalytic polypeptide (197–1594 bp). *GDG* exhibits a different architecture, encoding a major 480-amino acid polypeptide (105–1547 bp) and a shorter 57-amino acid chain (1593–1766 bp). Multiple sequence alignment with homologs from six Anacardiaceae species revealed distinct evolutionary conservation patterns among these genes (Fig. 3A). GDG exhibited the highest overall conservation, with 72% of residues showing ≥80% similarity across all species—notably concentrated in the C-terminal catalytic domain. Specifically, 346 of 480 positions in the GDG primary polypeptide displayed strong conservation, consistent with its role as the terminal committed enzyme directly controlling HT end-product structure. In contrast, PGM and BX9 displayed greater sequence divergence, with only 58% (182/312 residues) and 45% (209/465 residues) showing ≥80% conservation, respectively. Within the conserved regions, 142 positions in PGM and 128 positions in BX9 were completely identical across species, while the remaining conserved sites tolerated physicochemically similar substitutions.

**Figure 3.**
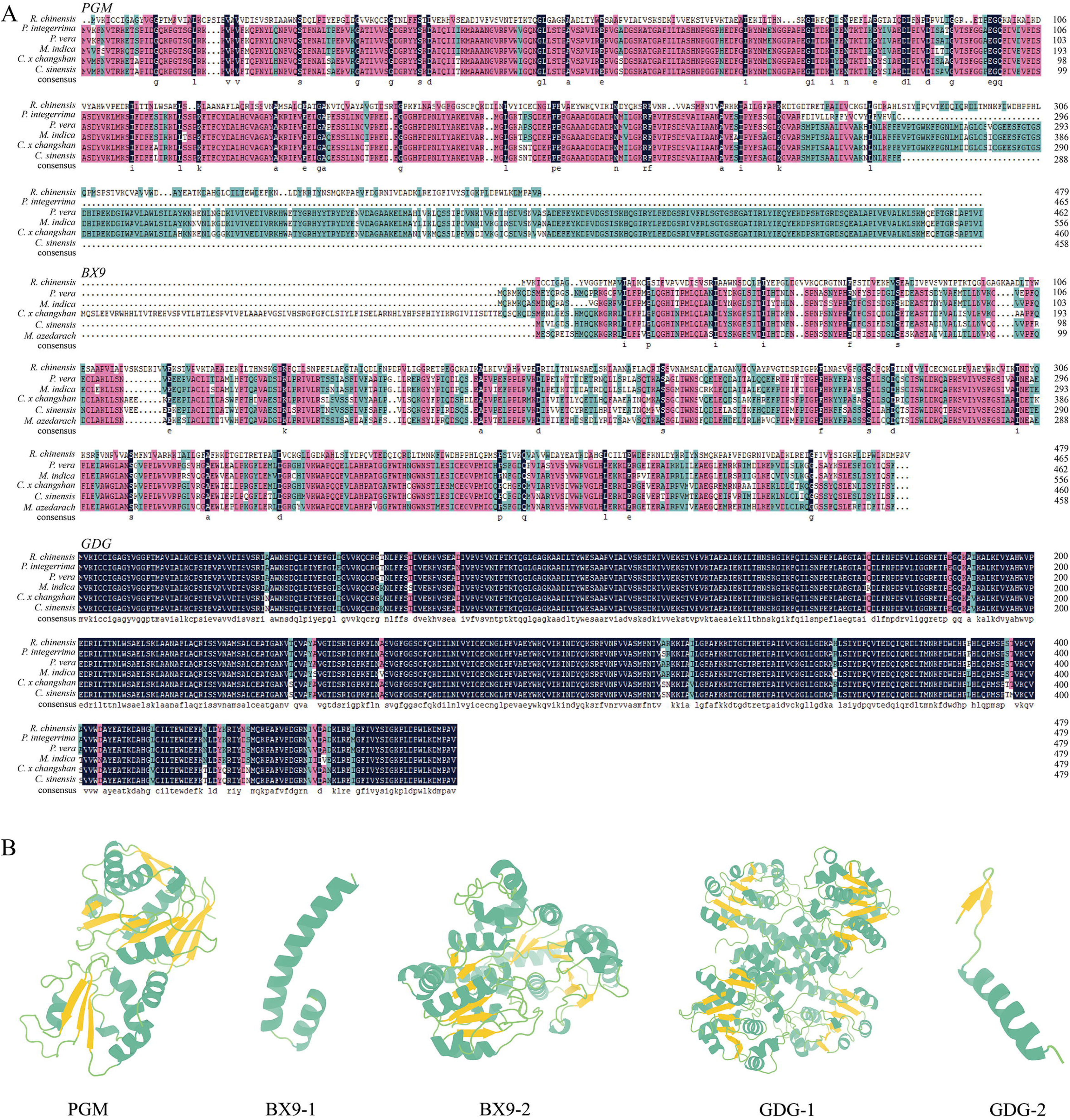
Molecular characterization and structural analysis of key genes in HT biosynthesis pathway. **(A)** Multiple sequence alignment of *PGM*, *BX9*, and *GDG* amino acid sequences with homologs from related Anacardiaceae species. Conservation levels are color-coded based on percentage conservation across all seven species: black background with white letters indicates completely conserved residues (100% identity) that are identical across all aligned sequences; dark pink/magenta background with white letters represents highly conserved sites (≥80% conservation) with strong similarity or conservative substitutions in at least five of six species; cyan/teal background with white letters denotes moderately conserved residues (60-79% conservation) sharing similar physicochemical properties in 4-5 of six species; white background with black letters indicates poorly conserved or variable positions (<60% conservation) with frequent amino acid substitutions; dots (…) represent alignment gaps or insertion/deletion events. Numbers on the right indicate amino acid positions. **(B)** Predicted three-dimensional protein structures of the encoded peptides. Structures are displayed in ribbon representation with α-helices shown in green and β-sheets in yellow.

The first peptide encoded by *PGM* is too short to predict its tertiary structure. The second PGM peptide (PGM) contains 103 atoms in α-helices, 62 atoms in β-sheets, and 147 atoms in loops. The first peptide of BX9 (BX9-1) comprises 40 atoms and 10 residues in α-helices, 4 atoms and 1 residue in loops, and no β-sheets or β-turns. The second BX9 peptide (BX9-2) consists of 221 atoms in α-helices, 57 atoms in β-sheets, and 179 atoms in loops. In a similar manner, the first peptide of GDG (GDG-1) comprises α-helices, β-sheets, and loops, with 439, 133, and 370 atoms, respectively. The shorter GDG peptide (GDG-2) consists of 17 atoms in α-helices, six atoms in β-sheets, and 8 atoms in loops (Fig. 3B, Table 1).

**Table 1.** Secondary structures of peptides.

| | | α-helices | β-sheets | β-turns | Loops |
| --- | --- | --- | --- | --- | --- |
| PGM | amino acid residues | 0 | 0 | 0 | 0 |
|  | atoms | 103 | 62 | 0 | 147 |
| BX9-1 | amino acid residues | 10 | 0 | 0 | 1 |
|  | atoms | 40 | 0 | 0 | 4 |
| BX9-2 | amino acid residues | 0 | 0 | 0 | 0 |
|  | atoms | 221 | 57 | 0 | 179 |
| GDG-1 | amino acid residues | 0 | 0 | 0 | 0 |
|  | atoms | 439 | 133 | 0 | 370 |
| GDG-2 | amino acid residues | 0 | 0 | 0 | 0 |
|  | atoms | 17 | 6 | 0 | 8 |

*PGM*, *BX9*, and *GDG* were individually or combinatorially overexpressed in *Arabidopsis thaliana* to validate their regulatory roles in plant HT biosynthesis (Fig. 4A, B). When expressed individually, *PGM* and *GDG* exhibited the highest expression levels; however, *BX9* showed peak expression in the *BX9*_*GDG* co-overexpression line, with overall higher expression observed in double overexpression lines (Fig. 4C–E). Protein expression levels varied across different transgenic lines. In single overexpression lines, *PGM* displayed the highest expression, followed by *GDG*. In double overexpression lines, target protein expression was consistently elevated, with particularly high levels detected in *PGM_GDG*-2, *PGM_BX9*-2, *PGM_BX9*-3, and *BX9_GDG*-1 plants (Fig. 5A).

**Figure 4.**
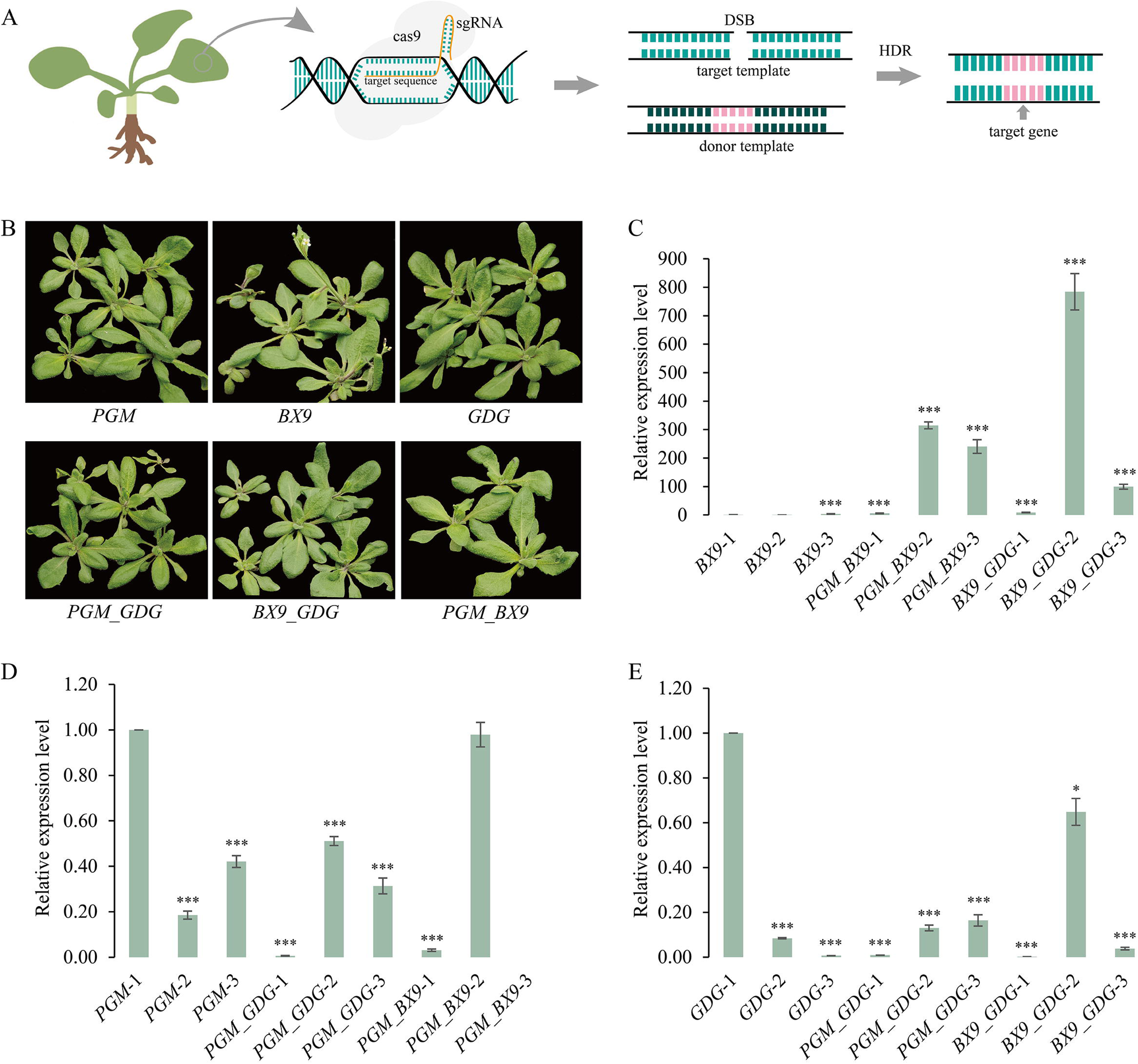
Functional validation of HT biosynthesis genes in transgenic Arabidopsis. (A) Gene overexpression strategy for single and combinatorial expression. (B) Representative transgenic lines. (C-E) RT-qPCR validation of transgene expression in (C) PGM, (D) BX9, and (E) GDG overexpression lines. * = P< 0.05; ** = P < 0.01; *** = P < 0.001.

**Figure 5.**
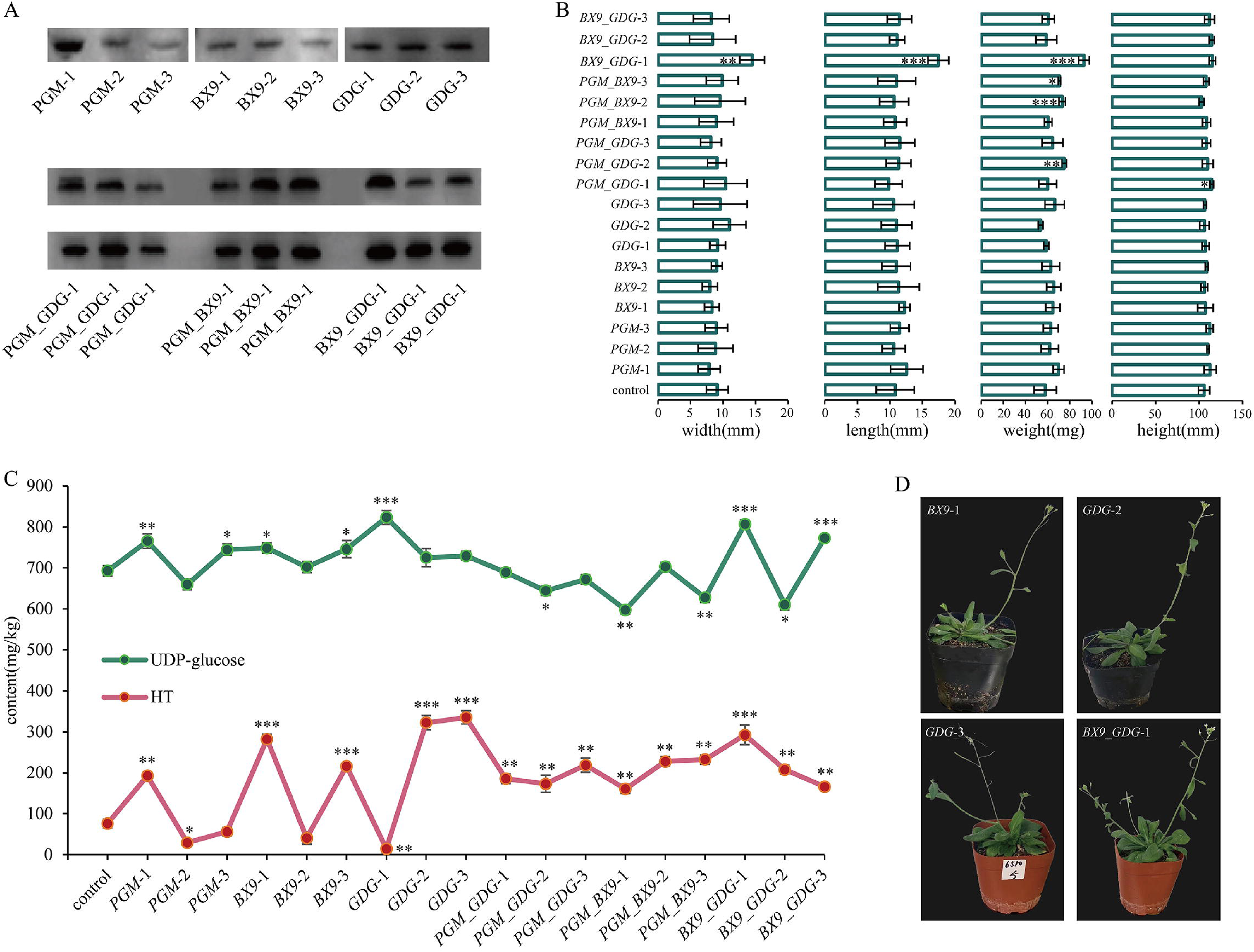
Protein expression and phenotypic characterization of transgenic *Arabidopsis* lines. **(A)** Western blot analysis of target protein expression in transgenic lines. **(B)** Morphological measurements of transgenic plants compared to wild-type control. **(C)** Biochemical quantification of UDP-glucose and HT content in transgenic lines. UDP-glucose content was altered in both single and combinatorial overexpression plants. HT content exhibited a consistent trend in single overexpressing lines, and in double overexpression lines, HT content was uniformly and significantly elevated regardless of gene combination. **(D)** Four transgenic lines with markedly enhanced tannin content: *BX9*-1, *GDG*-2, *GDG*-3, and *BX9*_*GDG*-1. These lines demonstrate the successful functional validation of the HT biosynthesis pathway genes in a heterologous plant system and their potential application in biotechnology. * = P < 0.05; ** = P < 0.01; *** = P < 0.001.

Among all treatment groups, only *BX9_GDG*-1 exhibited significantly greater leaf length and width compared to the control group (P < 0.05). The remaining transgenic lines demonstrated no significant differences in leaf dimensions. *PGM_GDG*-2, *PGM_BX9*-2, and *BX9_GDG*-1 plants exhibited a marked increase in fresh weight in comparison to the control, while only *PGM_GDG*-1 plants demonstrated a significant increase in plant height. No significant differences in weight or height were observed in the remaining transgenic lines relative to the control (Fig. 5B).

In the transgenic lines tested—both single and combinatorial overexpression plants targeting genes either upstream or downstream in the tannin biosynthesis pathway—UDP-glucose content was altered. Of the 18 tested lines, 6 showed no significant difference in UDP-glucose levels compared to the control, 4 exhibited significantly lower levels, and 7 displayed significantly higher levels, with the *BX9_GDG*-1 and *BX9_GDG*-3 lines showing the highest accumulation. The HT content exhibited a more consistent trend: in single overexpressing lines, one line each from the *PGM*, *BX9*, and *GDG* groups reduced HT levels relative to the control, while the remainder exhibited significantly increased HT accumulation, with two *GDG* lines showing over a threefold increase (Fig. 5C). In double overexpression lines, HT content was uniformly and significantly elevated regardless of gene combination. Four transgenic lines with markedly enhanced tannin content were ultimately identified: *BX9-1*, *GDG-2*, *GDG-3*, and *BX9_GDG-1* (Fig. 5D).

## Discussion

### Multi-gene Coordinated Regulation of Extreme HT Accumulation in Galls

The *S. chinensis*-*R. chinensis* system exemplifies one of nature’s most extreme cases of plant chemical defense, with gall HT concentrations reaching 74.49%—the highest documented in any plant-insect interaction and representing a 32-fold elevation over normal leaves. Our identification of the *PGM*-*BX9*-*GDG* regulatory module governing this remarkable biosynthetic capacity addresses the gap in understanding how insects manipulate host plant secondary metabolism to such extraordinary levels. The close correlation between gene expression dynamics and HT accumulation patterns strongly suggests aphid-derived effectors orchestrate a sophisticated molecular program intrinsic to the plant (Yang et al., 2024).

Central to this regulatory architecture is GDG, the terminal enzyme in the pathway, which exhibited the highest expression levels and temporal patterns mirroring HT dynamics. This suggests *GDG* functions as a rate-limiting switch controlling metabolic flux through the final biosynthetic step. Functional validation in *Arabidopsis* demonstrated that precise manipulation of this terminal enzyme produces more pronounced HT enhancement than upstream interventions, with double-gene transformants (particularly *BX9*_*GDG*) showing over threefold increases and more stable accumulation than single-gene approaches. This pattern aligns with metabolic control theory, which predicts flux control concentrates at committed steps rather than primary substrate provision (Noor & Liebermeister, 2024). The observation that UDP-glucose levels showed no positive correlation with final HT content further supports this framework—terminal pathway enzymes exert stronger control than upstream substrate availability.

The requirement for multi-gene coordination to achieve stable, high-level HT production mirrors recent discoveries in other plant systems. In strawberry, a "galloylation-degalloylation cycle" controlled by coordinated action of *FaUGT84A22*, *FaSCPL3-1*, and *FaCXE1/FaCXE3/FaCXE7* regulates HT homeostasis (Tahara et al., 2024). Similarly, in oak species, differential expression of *UGT84A13* and its transcriptional regulators *WRKY32*/*WRKY59* determines interspecific variation in tannic acid levels (Yang et al., 2024). These parallel findings suggest multi-gene regulatory networks may represent a conserved evolutionary strategy for fine-tuning HT biosynthesis across diverse plant lineages, enabling precise metabolic responses to environmental challenges including herbivory. Our demonstration that single-gene overexpression triggers compensatory negative feedback while multi-gene manipulation overcomes such regulation provides mechanistic insight: coordinated modulation of multiple pathway nodes may be essential to bypass homeostatic controls that normally constrain metabolite accumulation within physiologically tolerable ranges.

### Evolutionary Dynamics of Pathway Enzyme Conservation

Comparative sequence analysis revealed contrasting evolutionary patterns among the three regulatory genes. *PGM* and *BX9* show significant amino acid divergence from close relatives, potentially altering enzyme kinetics and substrate specificity through modifications in active site geometry (Judge et al., 2024). In contrast, *GDG* exhibits remarkable conservation across Anacardiaceae, underscoring its fundamental role in core HT synthesis. This pattern—wherein terminal biosynthetic enzymes display greater sequence conservation than upstream regulatory steps—parallels observations across diverse specialized metabolic pathways. In glucosinolate biosynthesis, terminal side-chain modification enzymes show higher interspecific conservation than upstream regulatory genes (Beekwilder et al., 2008; Kliebenstein et al., 2001), while in anthocyanin pathways, late biosynthetic genes (*DFR*, *ANS*, *UFGT*) are more conserved than early steps (*CHS*, *CHI*) (Rausher et al., 1999; Shimada et al., 2003). This evolutionary pattern likely reflects differential selection pressures: stronger purifying selection acts on enzymes catalyzing committed steps that directly determine end-product structure and biological activity, whereas enzymes serving multiple metabolic pathways tolerate greater sequence variation (Moghe & Last, 2015; Weng, 2014). The divergence in *PGM* and *BX9* may enable adaptive fine-tuning of pathway flux in response to specific ecological pressures, while *GDG* conservation ensures catalytic fidelity for the critical terminal reaction. Recent reconstitution studies of HT biosynthesis in *Nicotiana benthamiana* underscore the modular nature of this pathway and the feasibility of metabolic engineering approaches (Oda-Yamamizo et al., 2023; Tahara et al., 2024), with implications for enhancing plant defenses or producing HTs for pharmaceutical applications.

### Laccase-Mediated Detoxification: A Paradigm Shift in Understanding Tannin Adaptation

*S. chinensis* employs a laccase-dominated enzymatic system rather than tannase for HT detoxification challenges prevailing assumptions about insect adaptation to tannin-rich diets. Tannase activity remained negligible (0.08 nmol/min/g) throughout aphid development, while laccase activity peaked at 1649.86 nmol/min/g—representing a >20,000-fold difference in enzymatic capacity. This unequivocally demonstrates laccases, not tannases, constitute the primary HT-degrading system in this specialized herbivore. Three laccase genes (*Sc-Lac1*, *Sc-Lac5*, *Sc-Lac7*) were identified, with *Sc-Lac1* exhibiting highest expression and predominant abdominal localization, indicating gut-specific detoxification rather than salivary pre-digestion.

This compartmentalized enzyme distribution aligns with aphid feeding ecology: while stylet penetration establishes feeding sites on the inner gall wall, bulk HT degradation occurs post-ingestion in the digestive tract. Such spatial separation may serve multiple adaptive functions. First, concentrating laccase activity in the gut prevents premature oxidation of HT before ingestion, which could increase tannin-protein binding and reduce nutritional value (Barbehenn & Constabel, 2011). Second, it maximizes detoxification efficiency at the primary site of nutrient absorption, where HT toxicity would otherwise compromise gut epithelial integrity (Appel & Schultz, 1992). Third, it may facilitate conversion of degradation products into metabolically useful compounds—a hypothesis warranting future investigation through metabolomics approaches.

The efficacy of laccase-mediated degradation is demonstrated by *in vitro* assays showing 39–54% reduction in HT content following treatment with aphid homogenates. This confirms functional capacity for intestinal HT metabolism. Importantly, sexual aphids inhabiting moss (non-tannin environment) exhibit 13-fold lower laccase activity (127.47 vs. 1649.86 nmol/min/g), supporting facultative upregulation in response to dietary HT exposure rather than constitutive overexpression. This inducible strategy contrasts with constitutive detoxification observed in some generalist herbivores and may reflect a trade-off between maintaining high basal enzyme levels versus metabolic economy (Beran & Petschenka, 2022).

Laccase functional versatility explains this apparent dichotomy in biological roles. While plant laccases primarily polymerize lignin monomers and reinforce cell walls (Blaschek et al., 2023), insect laccases have diversified to serve cuticle hardening, wound repair, and—as our data demonstrate—dietary toxin degradation (Asemoloye et al., 2025; Sondhi et al., 2023). This functional divergence stems from the fundamental catalytic mechanism of laccases as multicopper oxidases generating free radicals; their ultimate function depends on cellular redox environment, substrate availability, and cofactor interactions (Jia et al., 2024). In the aphid gut’s reducing environment with abundant HT substrates, laccase activity channels toward oxidative degradation rather than polymerization, effectively neutralizing a potent plant defense. Recent discoveries of gut symbiont-mediated detoxification in other insect systems (Peterson, 2024; Rupawate et al., 2023) raise the question whether *S. chinensis* microbiome contributes to HT metabolism. While we demonstrate endogenous laccase capacity, symbionts with enhanced esterase and laccase activities have been documented in cypermethrin-degrading bacteria (Kline & Joshi, 2024). Future metagenomic analyses could elucidate potential synergistic interactions between host enzymes and microbial metabolism in achieving extreme tannin tolerance.

### Systemic HT Distribution and Gall Developmental Strategy

The uniform HT distribution between inner and outer gall walls throughout development provides mechanistic insights into gall biology. Despite concentrated aphid feeding on the inner wall, no spatial HT gradient was observed, indicating systemic rather than localized biosynthetic regulation. This homogeneity contrasts sharply with wound-induced defenses, which typically display steep concentration gradients radiating from damage sites (Vlot et al., 2021). The spatial uniformity reflects whole-organ metabolic reprogramming characteristic of gall systems, where insect effectors trigger developmental cascades affecting entire structures rather than localized responses (Desnitskiy et al., 2023; Gätjens-Boniche et al., 2023).

This systemic HT accumulation likely serves multifunctional defensive roles. Consistent high concentrations throughout gall tissues provide broad-spectrum chemical protection against diverse natural enemies, from fungal pathogens to parasitoid wasps and chewing herbivores (Kariñho-Betancourt et al., 2019). Recent transcriptomic analyses of oak galls demonstrate uniform upregulation of phenolic biosynthesis genes throughout gall tissues independent of feeding zone location (Betancourt et al., 2020), supporting the interpretation that galls function as insect-controlled plant structures optimized through extended phenotype manipulation (Labandeira, 2021; Murakami et al., 2021).

The mechanistic basis of gall induction remains incompletely understood despite decades of research. While chemical effectors including phytohormone mimics are implicated(Hirano et al., 2025; Yang et al., 2018), recent evidence suggests endosymbiotic bacteria may mediate plant cell transformation analogous to *Agrobacterium*-induced crown galls (Gätjens-Boniche et al., 2023). The complex morphogenesis and biochemical regulation exhibited by *R. chinensis* galls—characterized by precisely controlled HT biosynthesis, nutritive tissue differentiation, and structural elaboration—hints at sophisticated genetic manipulation potentially involving horizontal gene transfer from insect-associated microbes. This represents a frontier question for understanding gall induction mechanisms.

### Coevolutionary Arms Race and Metabolic Engineering Implications

Our findings illuminate a sophisticated chemical arms race wherein plants weaponize secondary metabolism to lethal extremes while herbivores evolve corresponding biochemical countermeasures. The 32-fold HT elevation in galls represents an escalation to concentrations that would be lethal to most insects, yet *S. chinensis* not only tolerates but thrives in this environment through specialized laccase machinery. This bidirectional escalation exemplifies reciprocal coevolutionary adaptation, where each organism’s fitness depends on overcoming the other’s innovations (Mathur et al., 2024).

The evolutionary dynamics likely involve repeated cycles of plant defense innovation and insect counter-adaptation. Gall-inducing insects demonstrate extreme host conservatism (Korneyev et al., 2024; Stone et al., 2009), suggesting that mastering a particular host’s chemical defenses creates strong barriers to host shifts—once an insect lineage evolves specialized detoxification for one plant’s defenses, switching to a new host with different chemistry may be prohibitively costly. The *S. chinensis*-*R. chinensis* association exemplifies such specialization, with the aphid’s laccase system exquisitely tuned to degrade the specific gallotannin structures produced by its host.

This study reveals the bidirectional molecular mechanisms enabling an extreme plant-insect chemical interaction. On the plant side, the *PGM-BX9-GDG* regulatory module governs unprecedented HT accumulation through coordinated multi-gene action, with the terminal enzyme GDG serving as a critical flux control point. On the insect side, a specialized, highly active laccase system—rather than conventional tannase—enables survival in this chemically extreme environment through gut-localized, inducible detoxification (Fig. 6). These findings demonstrate how coevolutionary arms races drive both plant defensive innovations and insect counter-adaptations to remarkable biochemical extremes, generating systems of exceptional sophistication. Beyond illuminating fundamental evolutionary processes, our results provide a genetic and enzymatic foundation for metabolic engineering efforts aimed at harnessing or disrupting this interaction for agricultural and pharmaceutical applications.

**Figure 6.**
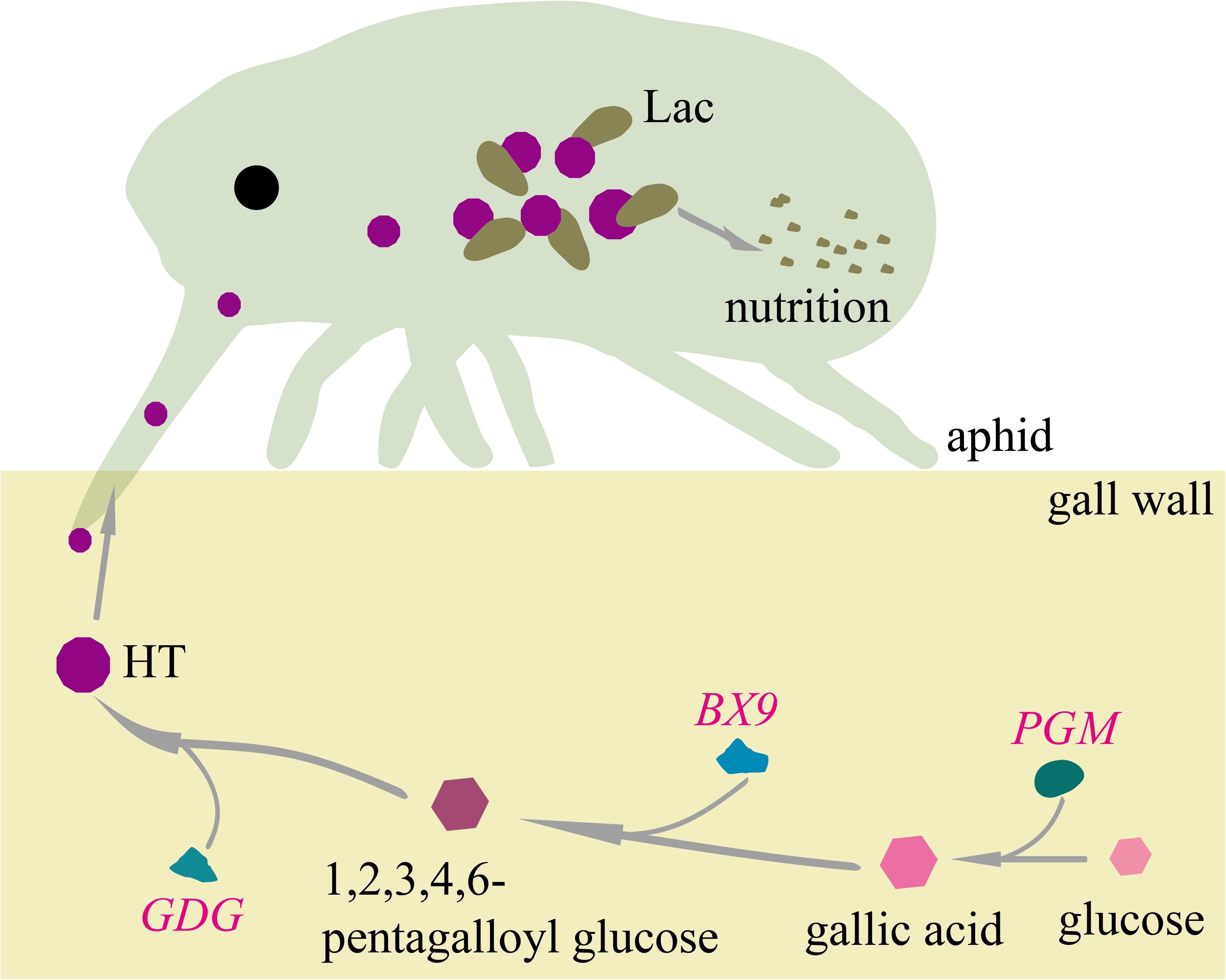
Molecular mechanisms of HT biosynthesis and degradation in aphid-gall interactions.

## Acknowledgments

This work was supported by grants from the Natural Science Foundation of Yunnan Province (202301AT070178), the National Natural Science Foundation of China (32370551) and the Expert workstation in Yunnan Province (202405AF140107).

## Author Contributions

HC conceived, planned and designed the research project. JL collected samples and performed sample preparation. QL, JL and WWW conducted the experiments. XZ performed microscopic imaging and photography. RH provided guidance on data analysis methods. QL and WWW analyzed and visualized the data. QL wrote the original draft of the manuscript. HC, GC and KKJ reviewed and edited the manuscript. HC and QL supervised the project and acquired funding. All authors have read and approved the final manuscript.

## Competing interests

We declare we have no conflict of interests.

## Data availability

The sequences of *PGM*, *BX9* and *GDG* have been deposited in the NCBI Genebank under accession PX688653, PX688654, PX688655.

## Notes

### Competing Interest Statement

The authors have declared no competing interest.

