## Supplementary material for "Molecular Arms Race: Tannin Biosynthesis and Laccase-Based Detoxification in the Aphid-Gall System": Table S1, Table S2, Table S3

**Table S1 Primers for RT-qPCR**

| Genes | Primers |
| --- | --- |
| *β-actin*-F3 | 5' - CTCTGTCTGGATTGGAGGGTC - 3' |
| *β-actin* -R3 | 5' - GGCTTGAGAAATGGTCGGA - 3' |
| *UDP*-F1 | 5' - AAGATGAAGCCTCCACCTCG - 3' |
| *UDP*-R1 | 5' - CAACATTTGACAGCAACTTAGCC - 3' |
| *PGM*-F1 | 5' - GGTGCTACACTTGTTGTTTCCG - 3' |
| *PGM*-R1 | 5' - TGACCAACCCATACTCGTCTTACT - 3' |
| *GDG*-F2 | 5' - CAATACCCCAACCAAGACACAG - 3' |
| *GDG*-R2 | 5' - TCCCTCAGCAAGAAACTCCG - 3' |
| *Sc-Lac1*-F | 5' - AGAAAATGTCGGTGGTTGGCAAGG - 3' |
| *Sc-Lac1*-R | 5' - CACCACCCATCAAAGCACTATAGC - 3' |
| *Sc-Lac5*-F | 5' - ATCCACCATCCGGTACTGTTACAG - 3' |
| *Sc-Lac5*-R | 5' - GGAACGAGTAGGTAACCATACCAC - 3' |
| *Sc-Lac7*-F | 5' - GTCATACACGAGCTACCACAACAC - 3' |
| *Sc-Lac7*-R | 5' - GAGACATCTTGTCGTCACCGCATT - 3' |
| *Sc-β-actin* -F | 5' - GGGGTGGATGAATTCATACGCACT - 3' |
| *Sc-β-actin* -R | 5' - AGAGCGCAAGTACTCTGTCTGGAT - 3' |

**Table S2 RACE adaptor primers**

| adaptor | Primers |
| --- | --- |
| 5’adaptor | GCTGTCAACGATACGCTACGTAACGGCATGACAGTGGGIIGGGIIGGGIIG |
| 5’adaptor | GCTGTCAACGATACGCTACGTAACGGCATGACAGTGTTTTTTTTTTTTTTTTTT |
| 5.3’outer | GCTGTCAACGATACGCTACGTAAC |
| 5.3’inner | GCTACGTAACGGCATGACAGTG |

**Table S3 RACE gene-specific primer** (GSP)

| Genes | Primers |
| --- | --- |
| *GDG* - F1 (3’) | ACAAGCTGAGGGAGATCGGTTTCATTG |
| *GDG* - F2 (3’) | GTAAGCCCCTGGATCCATGGCTGA |
| *GDG* - R1 (5’) | CCATAGTAGGCCCCCCAACATATCCA |
| *GDG* - R2 (5’) | CACCAATGCAGCAGATCTTCACCA |
| *GDG* - RT1 (5’) | ATTGCAGTTCCCTCAGCAAGA |
| *GDG* - RT2 (5’) | GTTTGAGAGGATTTGGAATTTGA |
| *PGM* - F5 (3’) | CAATTTTTGACTTCGAGTCTATTAAGAAGCTGCT |
| *PGM* - F6 (3’) | TTACATTCTGCTATGATGCACTTCATGGAGTT |
| *PGM* - R3 (5’) | TTGCGAGTGACGTTAAACTTCGCCAT |
| *PGM* - R4 (5’) | TTAGAAATTCTTTGAGGATGCTGACGAGAGA |
| *PGM* - RT1 (5’) | GGACCACCTGGATTGTGACTTG |
| *PGM* - RT2 (5’) | TTGCCTTGGATCCATCAGC |
| *BX9* - F1 (3’) | CTTTGGAAGCATTGTTGCTATTAATGAAACTG |
| *BX9* - F2 (3’) | CGTGGGGATTAGCCAATAGTGGGGT |
| *BX9* - R3 (5’) | TTCAAGTTGGTGTGAATTATGGTGATTGAAAG |
| *BX9* - R4 (5’) | TGTTTGCCAGCTGAAGCATGGGACT |
| *BX9* - RT1 (5’) | TGTAGAAGTCCTGGAATGGCTGT |
| *BX9* - RT2 (5’) | AAGCAATACTACTTGTTCTTAACACGAT |

**Table S4 CRISPR ORF amplification primers** (red sequences indicate vector recombination arms)

| Genes | Primers |
| --- | --- |
| *GDG*-6His | CZ-*GDG*-BglII-F: AACACGGGGGACTCTTGACCATGGTGAAGATCTGCTGC |
| *GDG*-6His | CZ-*GDG*-BstEII-R: GGAAATTCGAGCTGGTCACCCTAGTGGTGGTGGTGGTGGTGTGCCACTGCAGGCATGTC |
| *PGM*-6His | CZ-*PGM*-BglII-F: AACACGGGGGACTCTTGACCATGGCGAAGTTTAACGTCAC |
| *PGM*-6His | CZ-*PGM*-BstEII-R: GGAAATTCGAGCTGGTCACCCTAGTGGTGGTGGTGGTGGTGCCTTTTACCAAGGACCAT |
| *UDP*-3Flag | CZ-*UDP*-KpnI-F: GCGGGTCGACGGTACCATGCAGAAGATGGAGCAG |
| *UDP*-3Flag | CZ-*UDP*-KpnI-3flagR: CGTCATCCTTGTAATCGAACGATTGTATGTAACCAATC |
| *GDG*-6His+*UDP*-3Flag | CZ-*UDP*-KpnI-F: GCGGGTCGACGGTACCATGCAGAAGATGGAGCAG |
| *GDG*-6His+*UDP*-3Flag | CZ-3×FLAG-KpnI-R: TAGACATATGGGTACCTCATTTATCGTCATCATCTTTG |
| *PGM*-6His+*UDP*-3Flag | CZ-*UDP*-KpnI-F: GCGGGTCGACGGTACCATGCAGAAGATGGAGCAG |
| *PGM*-6His+*UDP*-3Flag | CZ-3×FLAG-KpnI-R: TAGACATATGGGTACCTCATTTATCGTCATCATCTTTG |
| *PGM*-6His+*GDG*-Flag | CZ-*GDG*-KpnI-F: GCGGGTCGACGGTACCATGGTGAAGATCTGCTGC |
| *PGM*-6His+*GDG*-Flag | CZ-*GDG*-KpnI-R: TAGACATATGGGTACCCTACTTGTCATCGTCGTCCTTGTAATCTGCCACTGCAGGCATGTC |
